# Scoping reviews in medical education: A scoping review

**DOI:** 10.1101/2020.07.23.218743

**Authors:** Lauren A. Maggio, Kelsey Larsen, Aliki Thomas, Joseph A. Costello, Anthony R. Artino

**Affiliations:** Uniformed Services University of the Health Sciences in Bethesda, Maryland, USA; School of Physical and Occupational Therapy and Associate Member, Institute of Health Sciences Education, Faculty of Medicine, McGill University, Montreal, Quebec, Canada; The George Washington University School of Medicine and Health Sciences, Washington, DC, USA

## Abstract

**Purpose:** The purpose of this study was to characterize the extent, range, and nature of scoping reviews published in core medical education journals. In so doing, the authors identify areas for improvement in the conduct and reporting of scoping reviews, and highlight opportunities for future research.

**Method:** The authors searched PubMed for scoping reviews published between 1999 through April 2020 in 14 medical education journals. From each review, the authors extracted and summarized key bibliometric data, the rationales given for conducting a scoping review, the research questions, and key reporting elements as described in the PRISMA-ScR reporting guidelines. Rationales and research questions were mapped to the reasons for conducting a scoping review, as described by Arksey and O’Malley.

**Results:** One hundred and one scoping reviews were included. On average 10.1 scoping reviews (MED=4, SD=13.08) were published annually with the most reviews published in 2019 (n=42) in 13 of the included 14 journals reviewed. *Academic Medicine* published the most scoping reviews (n=28) overall. Authors described multiple reasons for undertaking scoping reviews, including to: summarize and disseminate research findings (n=77); examine the extent, range, and nature of research activity in a given area (n=74); and to analyze an emerging topic or heterogenous literature base (n=46). In 11 reviews there was alignment between the rationales for the scoping review and the stated research questions. No review addressed all elements of the PRISMA-ScR, with only a minority of authors publishing a protocol (n=2) or including stakeholders (n=20). Authors identified several shortcomings of scoping review methodology, including being unable to critically assess the included studies.

**Conclusions:** Medical educators are increasingly conducting scoping reviews with a desire to characterize the literature on a topic. There is room for improvement in the reporting of scoping reviews, including the alignment of research questions, the creation and publishing of protocols, and the inclusion of external stakeholders in published works.

## Introduction

Over the last two decades the number of knowledge syntheses published in core medical education journals has increased by 2,620%.^1^ Amongst these knowledge syntheses, there has been an even steeper rise in the number of scoping reviews published, with that number increasing by 4,200%. Additionally, medical education scholars have recently highlighted various aspects of scoping reviews in articles,^2–4^ a book chapter,^5^ and a podcast.^6^ The growth of scoping reviews and related scholarly attention suggests they may be playing an increasingly important role in the field’s discourse and, as such, have potential to positively influence future research and education policy. However, despite this growth and attention, the extant research on scoping reviews provides limited information about the nature of the scoping reviews in medical education, including how they are conducted or why medical educators decide to undertake this type of knowledge synthesis. This lack of direct insight about scoping reviews makes it difficult to know where the field stands and may hamper attempts to take evidence-informed steps to improve the conduct, reporting, and utility of scoping reviews in medical education.

Scoping reviews are often cast as publications that “map” the depth and breadth of the literature in a field.^7,8^ Through such synthetic mapping, authors describe the main concepts that underpin a topic and can illuminate gaps in the literature. Scoping reviews are generally driven by broad, exploratory research questions and typically incorporate studies that employ a variety of research designs.^9,10^ In their seminal article outlining a model for scoping reviews, Arksey and O’Malley described a six-step framework for conducting scoping reviews.^7^ These steps include: (1) identifying the research question, (2) identifying relevant studies, (3) selecting the studies to be included, (4) charting the data, (5) collating, summarizing, and reporting results, and (6) consultation with stakeholders. Step six, consultation with stakeholders, is an optional step in the original framework. Over time, scholars have suggested modifications to the steps.^9–12^ Some of these modifications are captured in the Preferred Reporting Items for Systematic Reviews and Meta-analysis Extension for Scoping Reviews (PRISMA-ScR),^13^ the first reporting guideline specific to scoping reviews.

Similar to the trends in medical education, the number of scoping reviews published in the health sciences is also on the rise.^14,15^ To characterize these scoping reviews, researchers have recently authored discipline-specific^16^ and cross-disciplinary^8,17^ scoping reviews of scoping reviews. Collectively, these scoping reviews have identified methodological shortcomings and a need for improved scoping review reporting. While these studies are valuable, two are now several years old and the most recent review focuses solely on rehabilitation medicine, thus providing limited information on current approaches specific to medical education. As a relatively new field that includes researchers from a variety of backgrounds and research traditions, we believe there is value in specifically examining scoping reviews in medical education. Doing so will help the field identify areas for improvement in the conduct and reporting of scoping reviews, thereby helping to ensure that those produced are relevant to and practical for application in the field (e.g., useful to map the literature of a topic, identify gaps in the literature). Thus, in this study, we aim to characterize the extent, range, and nature of scoping reviews published in medical education journals

## Methods

Guided by the framework presented by Arksey and O’Malley^7^ as updated by Levac,10 we conducted a scoping review of medical education scoping reviews to examine and characterize the extent, range, and nature of scoping reviews in core medical education journals.

### Identifying the research question

This scoping review is one part of a larger bibliometric analysis conducted by members of the author team. In the larger analysis, we broadly characterized knowledge syntheses in a core set of medical education journals.^1^ In that bibliometric analysis, we observed exponential growth in the number of scoping reviews. We argue that such growth likely has had a significant impact on the field’s ongoing scholarly discourse and future directions. This observation prompted three follow-on questions: (1) What are the characteristics of the scoping reviews?; (2) What rationales do authors provide for undertaking a scoping review?; and (3) How do authors report the details of their reviews?

### Identifying relevant studies

In the present study, we identified scoping reviews published during the time frame of the original study (1999-2019)^1^ plus those reviews published in the first four months of 2020. On March 26, 2020 JC, an information scientist, queried PubMed using a combination of keywords and controlled vocabulary terms (See Maggio 2020 for complete searches)^1^. JC reran this search on April 27, 2020 to capture any new citations. All retrieved citations and their metadata (e.g., abstract text, author names) were downloaded and managed in GoogleSheets.^18^ On May 21, 2020, JC obtained from Web of Science the number of times each review had been cited.

Searches were limited to 14 journals previously identified as core medical education titles (Lee, 2013; Maggio, 2018), including: *Academic Medicine, Advances in Health Sciences Education, BMC Medical Education, Canadian Medical Education Journal, Clinical Teacher, International Journal of Medical Education, Advances in Medical Education and Practice, Journal of Graduate Medical Education, Medical Education, Medical Education Online, Medical Teacher, Perspectives on Medical Education, Teaching and Learning in Medicine*, and *The Journal of Continuing Education in the Health Professions*.

To conduct the review, we carefully assembled a research team with expertise in knowledge synthesis methodology (AT, LM), information science (JC, LM), and medical education (LM, KL, AT, JC, AA) to guide the overall conduct of the review and our interpretation of the results. All team members had actively participated in the conduct of previous scoping reviews.

### Selecting the studies to be included

To select studies for inclusion, we used an iterative approach. LM and JC independently reviewed the titles and abstracts of all citations and met three times during the process to discuss coding discrepancies. AA was available to facilitate any coding disagreements. Articles were included if they described the conduct of a scoping review. Articles in which authors discussed scoping reviews as a methodological approach, but that did not describe undertaking an actual scoping review, were excluded.

### Charting the data

We created a data extraction tool that included and expanded upon the 22-items from the PRISMA-ScR checklist.^13^ The extraction tool also included questions about the review’s population, the authors’ rationale for undertaking the review, the stated research questions or aims, descriptions of the author team’s expertise, and any limitations of the scoping review approach identified by the authors (See https://doi.org/10.6084/m9.figshare.12699698.v1 for the data charting tool completed for each included study).

JC and LM piloted the data extraction tool by independently reviewing seven reviews and then comparing results. The extraction tool was modified based on the pilot and then used to extract data from the remaining full-texts of articles. JC and LM independently extracted data from all included articles and met three times to discuss any discrepancies. AA was again available to discuss any coding disagreements and serve as a tiebreaker.

### Collating, summarizing and reporting results

Descriptive statistics were calculated using GoogleSheets to describe characteristics of the scoping reviews (e.g., number of reviews published, author team makeup, items reported according to the PRISMA-ScR).

To describe the authors’ rationales for undertaking a scoping review and their research questions, we conducted a thematic analysis.^19^ To begin, JC and LM familiarized themselves with the data through multiple readings. During this process, they identified the relevance of the four reasons proposed by Arksey and O’Malley for conducting a scoping review. These reasons include: (1) to examine the extent, range, and nature of research activity in a given area; (2) to determine the value of undertaking a full systematic review; (3) to summarize and disseminate research findings; and (4) to identify gaps in the existing body of literature.^7^ Based on a whole-team call, we decided to use these four reasons as *a priori* codes while remaining open to additional rationales for preliminary coding. LM and JC independently coded all rationales and research questions and then met with KL. The meeting focused on cross-checking agreement on the overall coding of rationales and research questions and resolving any disagreements through discussion.

### Undertaking consultation

We shared our preliminary findings with seven stakeholders to understand if and in what ways our findings resonated with their experiences conducting scoping reviews. Stakeholders were authors of scoping reviews (n=6), editors of medical education journals (n=2), and faculty members in health professions education graduate programs (n=2). Stakeholders were asked to review our results and to weigh in on topics for discussion. All seven stakeholders agreed that our findings corresponded with their experiences and five of them provided suggestions for interpretation, which we tried to incorporate into our discussion.

## Results

In our initial bibliometric analysis, we had identified 88 scoping reviews plus one that was referred to as a systematic scoping review.^1^ The updated search and application of inclusion criteria resulted in 12 additional scoping reviews for a total of 101 included studies (Fig. 1).

**Figure 1:**
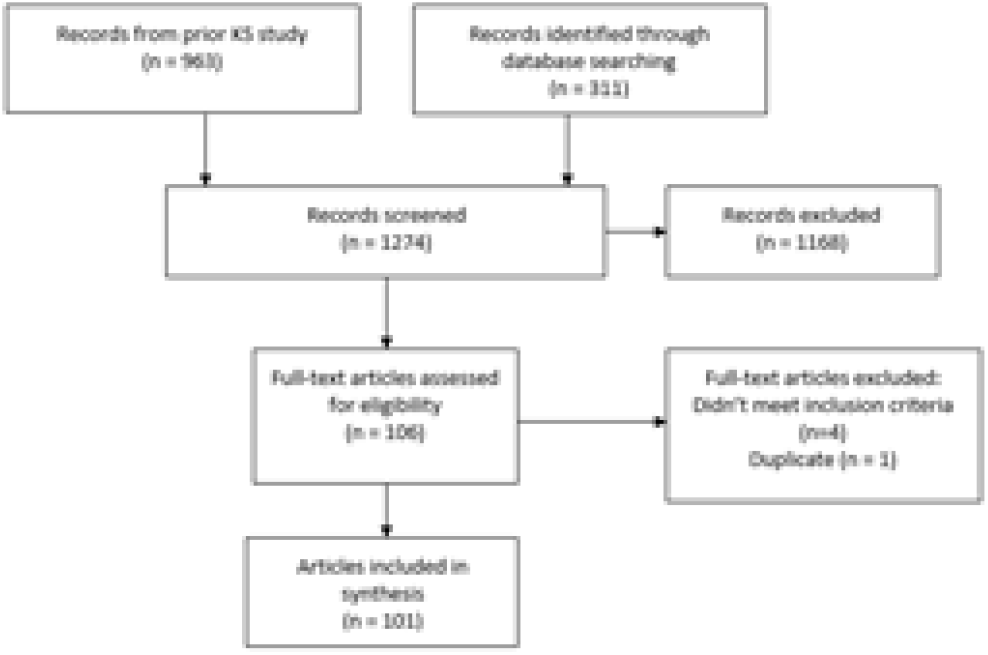
Flow Diagram of study inclusion process

On average 10.1 scoping reviews (SD=13.08, MED=4, Range 0-42) were published annually (Figure 2) with the most published in 2019 (n=42; 41.6%). The first scoping review in our sample was published in 2011. Scoping reviews were featured in 13 of the 14 journals with *Academic Medicine* (n=28, 27.7%), *Medical Education* (n=18, 17.8%), and *BMC Medical Education* (n=16, 15.8%) publishing the most. *Clinical Teacher* did not publish any scoping reviews during this time period. Thirty-nine scoping reviews (38.6%) reported receiving funding. All reviews synthesized journal articles, with 28.7% (n=29) also including book chapters, dissertations, websites, posters, and conference proceedings. Of those that focused only on journal articles, multiple reviews limited inclusion to empirical studies, thereby excluding commentaries, letters, editorials, and review articles.

**Figure 2:**
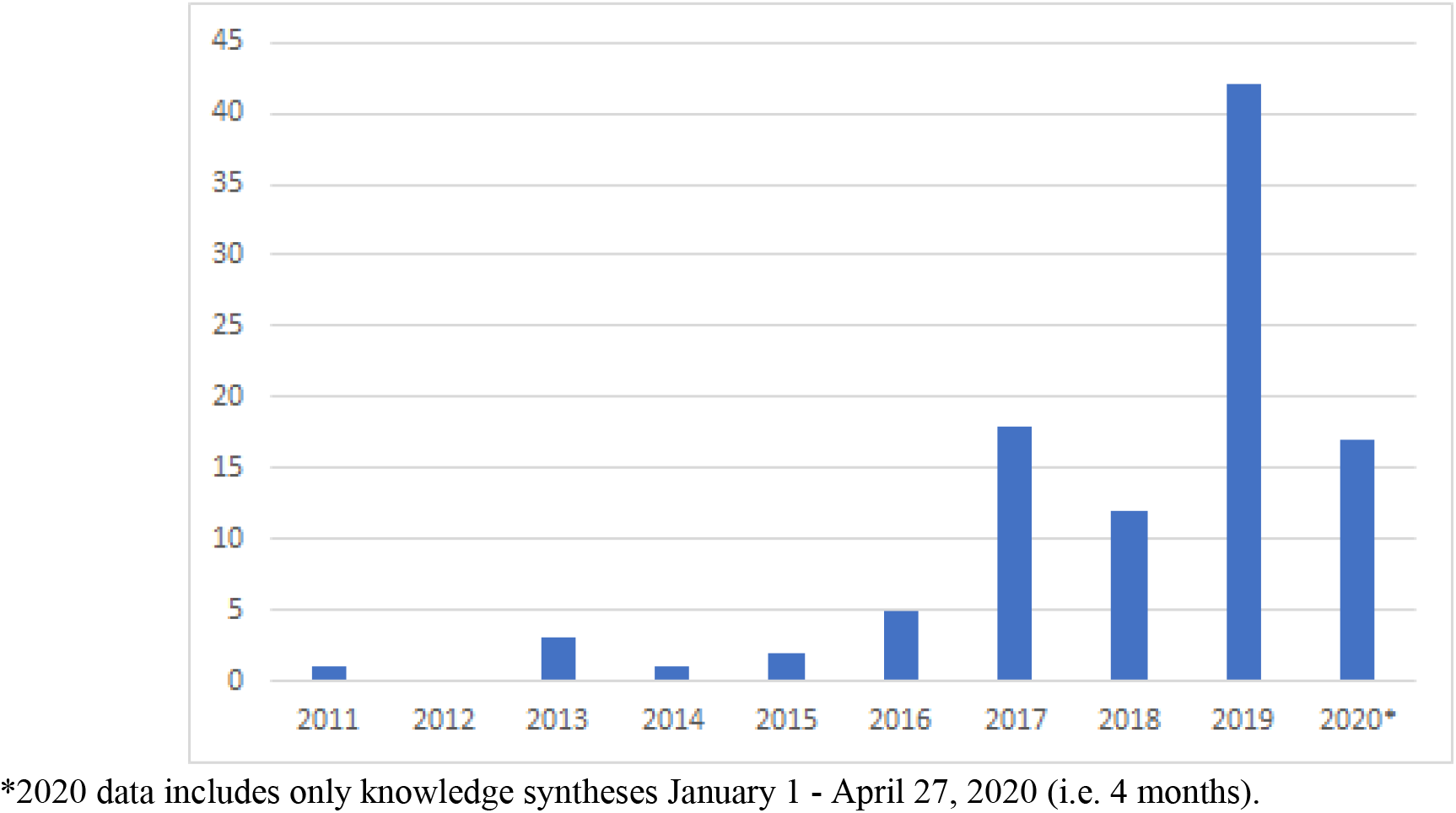
Counts of knowledge syntheses published between 2011 and April 27, 2020

Scoping reviews were authored, on average, by 5.33 authors per review (SD =2.92, MED=5, Range 1-17). Seventy-two reviews featured author teams from multiple institutions (71.3%) and multiple countries (n=25, 24.8%). A single review featured one author only.^20^ Lead authors were based in 16 countries with the majority in Canada (n=31, 30.7%), the United States (n=27, 26.7%), and Australia (n=9, 8.9%). In 23 reviews (22.8%), authors described their team members’ backgrounds and expertise in relation to their conduct of the scoping review. For example, authors of a scoping review on parenthood in residency noted that they “had knowledge and experience of parenthood during GME (JAB, SWS), literature reviews (ALB, KEE, SM), and information management and retrieval (ALB)”.^21^ Eight studies (7.9%) were led by doctoral students in health professions education.^22–29^

Scoping reviews with available citation data (n=89) were cited, on average, 6.37 times (SD=11.67, MED=2, Range 0-61). Eighteen articles (17.8%) had not been cited; of those, 12 (66.7%) were published after 2019. (See Table 1 for the top 10 most cited scoping reviews)

**Table 1:**
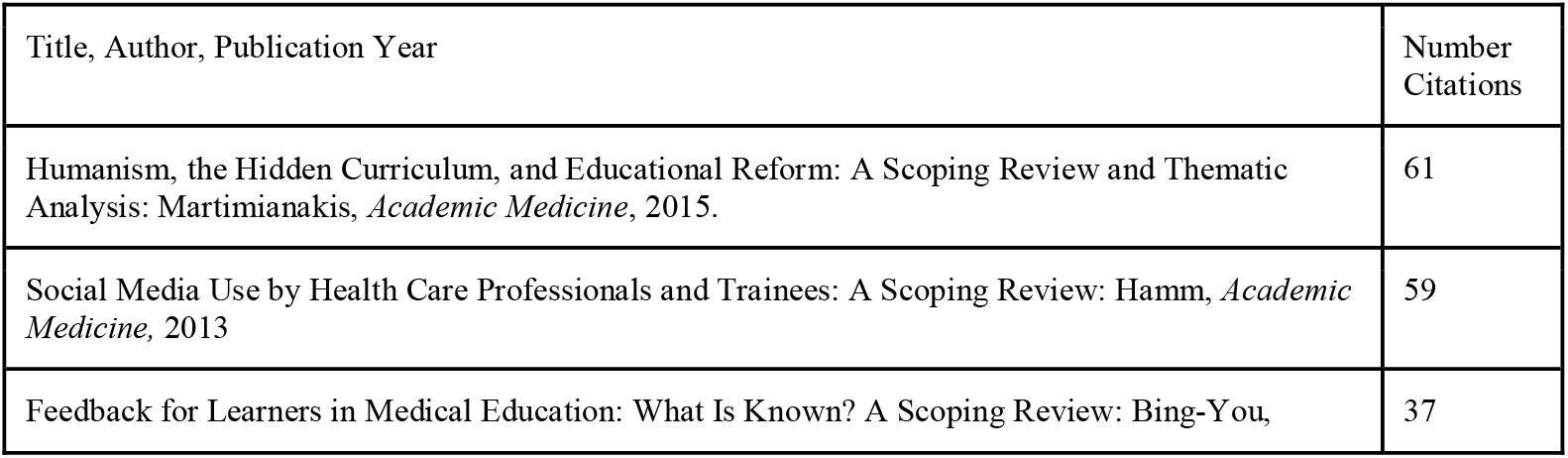
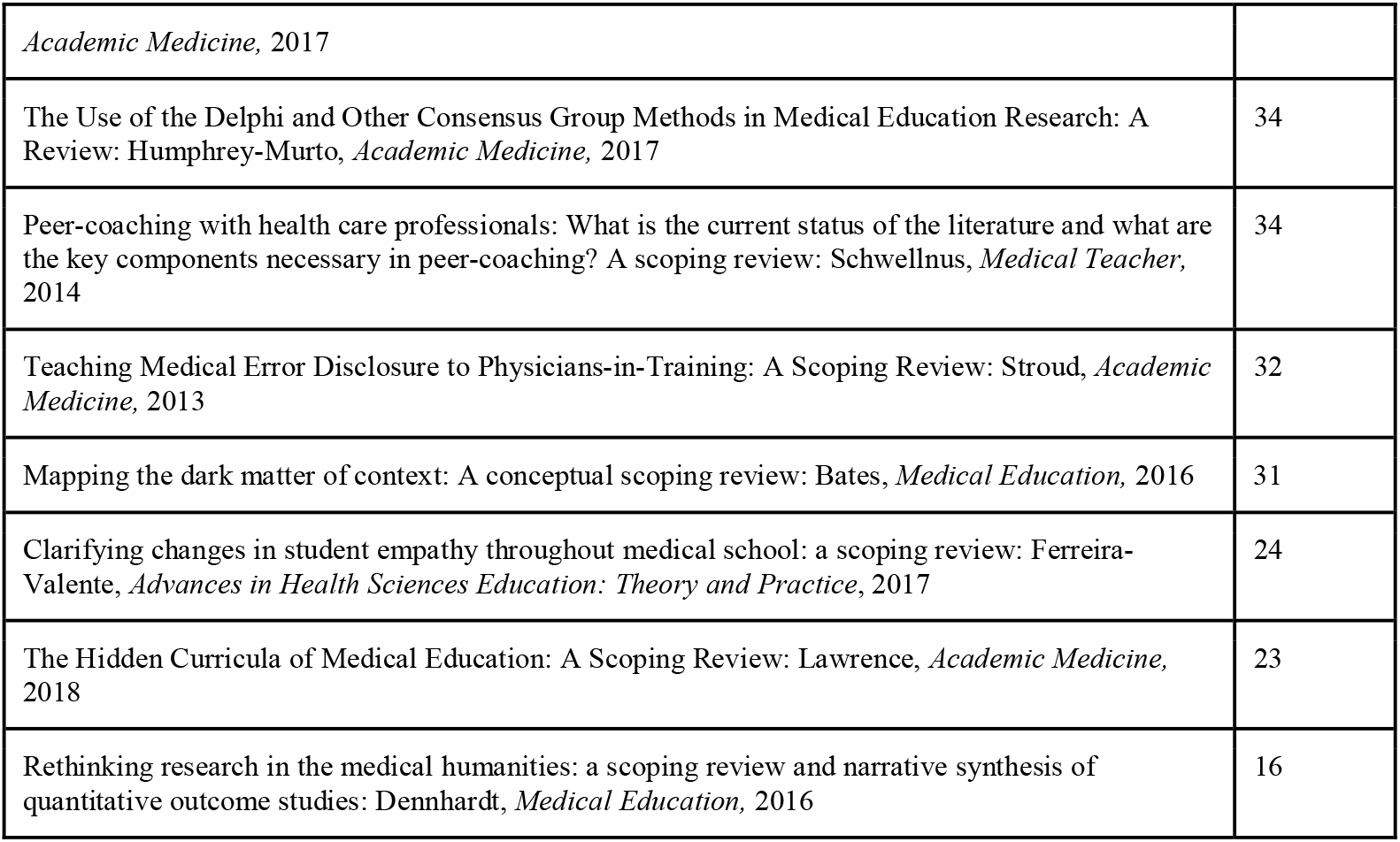
Top 10 Cited Scoping Reviews published in core medical education journals between 2009-April 2020

| Title, Author, Publication Year | Number Citations |
| --- | --- |
| Humanism, the Hidden Curriculum, and Educational Reform: A Scoping Review and Thematic Analysis: Martimianakis, *Academic Medicine*, 2015. | 61 |
| Social Media Use by Health Care Professionals and Trainees: A Scoping Review: Hamm, *Academic Medicine*, 2013 | 59 |
| Feedback for Learners in Medical Education: What Is Known? A Scoping Review: Bing-You, | 37 |
| *Academic Medicine*, 2017 |  |
| The Use of the Delphi and Other Consensus Group Methods in Medical Education Research: A Review: Humphrey-Murto, *Academic Medicine*, 2017 | 34 |
| Peer-coaching with health care professionals: What is the current status of the literature and what are the key components necessary in peer-coaching? A scoping review: Schwellnus, *Medical Teacher*, 2014 | 34 |
| Teaching Medical Error Disclosure to Physicians-in-Training: A Scoping Review: Stroud, *Academic Medicine*, 2013 | 32 |
| Mapping the dark matter of context: A conceptual scoping review: Bates, *Medical Education*, 2016 | 31 |
| Clarifying changes in student empathy throughout medical school: a scoping review: Ferreira-Valente, *Advances in Health Sciences Education: Theory and Practice*, 2017 | 24 |
| The Hidden Curricula of Medical Education: A Scoping Review: Lawrence, *Academic Medicine*, 2018 | 23 |
| Rethinking research in the medical humanities: a scoping review and narrative synthesis of quantitative outcome studies: Dennhardt, *Medical Education*, 2016 | 16 |

### Rationale for scoping reviews

Eighty-eight (87.1%) authors described rationales for selecting a scoping review methodology, with most referencing multiple rationales (n=82; 93.1%). There were, on average, 2.64 rationales stated per review (MED=3; SD=1.39). For example, one author described, “Literature regarding feedback for learners is far-ranging, has not been broadly assessed, contains varied approaches, and is not obviously suitable for systematic study. Therefore, we conducted a scoping review of the published literature to (1) identify the extent, range, and quantity of evidence available regarding feedback for learners in medical education; (2) map key concepts from the literature on feedback; and (3) determine existing gaps that may spur future research”.^30^

The most often stated rationales were: to summarize and disseminate research findings (n=77; 87.5%); to examine the extent, range, and nature of research activity in a given area (n=74; 84.1%); and to contend with the nature of the study topic or available literature (n=46; 52.3%). Of those who said they selected a scoping review methodology to contend with the nature of the topic or literature, 28 (31.8%) mentioned the heterogeneity of available literature, 15 (17.0%) said the topic was emerging, and eight (9.1%) noted both reasons. See Table 2 for all rationales.

**Table 2:** Rationales for conducting a scoping review and research questions in 101 scoping reviews published in 14 core medical education journals

| Rationales | For selecting scoping review methodology n=88 n (%) | Research question and/or study aims n=98 n (%) |
| --- | --- | --- |
| Summarize and disseminate research findings* | 77 (87.5) | 89 (90.1) |
| Examine the extent, range, and nature of research activity in a given area* | 74 (84.1) | 86 (87.8) |
| Contend with the nature of the study topic or available literature | 46 (52.3) | 18 (18.4) |
| Identify gaps in the existing body of literature* | 24 (27.3) | 13 (13.3) |
| Refine subsequent research inquiries (not including systematic reviews) | 15 (17.0) | 11 (11.2) |
| Demonstrate the use of theory or model implementation | 12 (13.6) | 7 (7.1) |
| Create future products (e.g., a framework, curriculum, policy) | 8 (9.1) | 8 (8.2) |
| Determine the value of undertaking a full systematic review* | 5 (5.7) | 1 (1.1) |
| Assess quality or effectiveness | 5 (5.7) | 5 (5.7) |

\*These rationales were described by Arksey and O'Malley.<sup>7</sup>

### Research Questions/Study aims

Ninety-eight authors (97.0%) included research questions and/or aims for their reviews. Authors put forth, on average, 2.40 research questions or aims per review (SD=.99, MED=2, Range 0-5). Similar to their rationales for conducting a scoping review, authors’ research questions or aims were attempting to: summarize and disseminate research findings (n=89; 90.8%); examine the extent, range, and nature of research activity in a given area (n=86; 87.8%); and contend with the nature of the study topic or available literature (n=18; 18.4%).

Although authors usually provided more rationales for selecting the scoping review methodology than they offered research questions, there was some alignment in our coding of the authors’ rationales and their research questions/aims. In 11 reviews (10.9%), the rationales for conducting a scoping review and the research questions were in complete alignment, such that we coded each in the exact same way. In 63 studies (62.4%) there was overlap such that the research questions indicated a desire to summarize the literature and examine its nature, but the rationale for selecting a scoping review included additional rationales, such as going further to describe the need to deal with heterogeneity of the available literature. For example, in one study the authors described their decision to undertake a scoping review to identify gaps in the research and clarify key concepts, as well as to clarify definitions of the concept; this aligned with, but went beyond, their research question, which was to describe the scope of the literature on the topic.^23^

### Reporting in alignment with PRISMA-ScR

Studies reported items from the PRISMA-ScR to varying degrees, and none included all of the items. Thirteen reviews (12.9%) cited following the PRISMA,^31^ and five (5.0%) the PRISMA-ScR.^13,32–36^ Table 3 summarizes the components of the PRISMA-ScR present in the included scoping reviews. For details by study, see https://doi.org/10.6084/m9.figshare.12699698.v1.

**Table 3:** A summary of the presence of the PRISMA-ScR Checklist^13^ items in 101 scoping reviews in 14 core medical education journals

| Checklist item | Number scoping reviews (%) |
| --- | --- |
| 1. Study identified as a scoping review in the title | 89 (88.1) |
| 2. Includes a structured abstract | 100 (99.0) |
| 3. Describes the rationale for the review in the context of what is already known<br>3.1 Explains why review questions/aims lend themselves to a scoping review approach | 88 (87.0)<br>98 (97.0) |
| 4. Provides the questions and objectives being addressed | 101 (100) |
| 5. Indicates whether a review protocol exists | 2 (2.0) |
| 6. Specifies characteristics of the sources of evidence used as eligibility criteria and provides rationale | 98 (97.0) |
| 7. Presents all information sources searched*<br>7.2 Includes the most recent search date | 101 (100)<br>50 (50.0) |
| 8. Includes the full search strategy for at least one database such that it could be repeated<br>8.1 Describes limits used | 61 (61.0)<br>100 (99.0) |
| 9. States the process for selecting evidence included | 100 (99.0) |
| 10. Describes the methods of charting data | 65 (64.3) |
| 11. Lists and defines all variables for which data was sought and any simplifications made | 75 (74.3) |
| 12. If done, provides a rationale for conducting critical appraisal of included sources | 13 (12.8) |
| 13. Described the methods of handling and summarizing the charted data | 86 (85.6) |
| 14. Provides the number of sources of evidence screened, assessed for eligibility, and included in the review ideally presented as a flow diagram | 97 (96.0) |
| 15. For each evidence source, presents characteristics for which data were charted and provide citations | 58 (57.4) |
| 16. If done, presents results of critical appraisal | 10 (9.9) |
| 17. For each included evidence source, present relevant data that were charted that related to the review questions and objectives | 70 (69.3) |
| 18. Summarizes charting results as they relate to research questions and objectives | 97 (96.0) |
| 19. Summarizes the main results, linking to review questions and objectives | 101 (100) |
| 20. Discusses the limitations of the scoping review process | 23 (22.8) |
| 21. Provides a general interpretation of the results with respect to the research questions/objectives | 101 (100) |
| 22. Describes funding sources of funding for the included sources of evidence | 0 (0) |
| 22. Describes the role of the funders of the scoping review | 39 (38.6) |

While the PRISMA-ScR is quite detailed, we charted additional study details based on Arksey and O’Malley’s framework as modified by Levac (See Table 4).^7,10^ Most reviews described following Arksey and O’Malley’s framework (n=73, 72.3%)^7^ and 34 (33.7%) of these reviews used Levac’s revision.^10^

**Table 4:** Summary of scoping review characteristics charted based on Arksey and O’Malley’s framework as modified by Levac^7^

| Identifying relevant studies | Number studies (%) |
| --- | --- |
| Conducted database searches<br>using multiple databases | 101 (100)<br>94 (93.1) |
| Handsearched included articles | 72 (71.3) |
| Included a librarian | 62 (61.3) |
| Consulted article authors | 4 (3.9) |
| **Selecting studies to be included** |  |
| Authors included physicians<br>Undergraduate Medical Education<br>Graduate Medical Education<br>Continuing Medical Education<br>Across all three levels | 55 (54.5)<br>37 (36.6)*<br>31 (30.7)*<br>21 (20.8)*<br>13 (12.9) |
| Authors included multiple health professions | 44 (43.6) |
| Authors included physiotherapists only | 2 (1.9) |
| Authors included emergency medical technicians only | 2 (1.9) |
| Authors included nurses only | 1 (.9) |
| **Data Charting** |  |
| Authors published data charting tool | 36 (35.6) |
| Authors piloted their data charting tool | 36 (35.6) |
| Data charting was done by more than one author | 65 (64.3) |
| **Collating, summarizing and reporting results** |  |
| Critical appraisal conducted<br>using Medical Education Research Quality Instrument | 12 (11.8)<br>10 (9.9) |
| Qualitative analysis conducted<br>Using thematic analysis<br>Using content analysis<br>Using narrative analysis | 61 (60.3)<br>38 (37.6)<br>10 (9.9)<br>5 (4.9) |
| Authors reported overall limitations | 94(93.0) |
| Authors reported limitations of scoping review methodology<br>Inability to conduct critical appraisal | 34 (33.6)<br>15 (14.8) |
| Consultations with key stakeholders |  |
| Authors reached out to key stakeholders | 20(19.8) |

\*Several studies included combinations of learner levels thus counts sum to greater than 55 studies.

## Discussion

Scoping reviews are increasingly conducted in medical education and are published by almost all of the core journals. These findings suggest that scoping reviews have the potential to play a powerful role in research and practice of medical education. As 38.6% of authors reported receiving funding for their work, with nearly half receiving taxpayer funds, it is critical that scoping reviews are well conducted and clearly reported. With this in mind, we focus our discussion on areas we feel are ripe for improvement in the conduct of scoping reviews.

Researchers have highlighted the importance of authors linking their rationale to their research questions. Doing so helps to guide the scoping review’s overall conduct, especially to inform the inclusion/exclusion of evidence and data extraction processes.^4,10^ We observed some alignment in authors’ rationales and their research questions/aims, but also noted some areas for improvement. For example, the majority of authors’ rationales and research questions mapped to those described by Arksey and O’Malley in 2005.^7^ In 2010, Levac criticized these rationales as being applicable broadly to a variety of knowledge synthesis methodologies and not necessarily specific to scoping reviews.^10^ Thus, it is possible that this lack of specificity contributed to the suboptimal alignment between the reported rationales and research questions. In light of this finding, we encourage medical education researchers to consider if Arksey and O’Malley’s rationales are really “fit for purpose” for the field of medical education. Additionally, the field might consider asking scoping review authors to specifically describe *why* they selected a scoping review methodology and what elements factored into consideration (e.g., the nature of the medical education literature, the intricacies of their topic, the contribution to the field, the expertise of their research team, and/or their personal needs such as a graduate student familiarizing herself with a topic). More clearly articulating medical education focused rationales could help scholars in the field determine if a scoping review is truly the right approach for a particular knowledge synthesis and help make explicit the unique contribution of this review type.

Author teams featured researchers from diverse backgrounds and multiple professions. Most teams included individuals with varied methodological training. This diversity can fundamentally impact, both positively and negatively, the conduct and reporting of a scoping review.^4^ The same can be said for relatively homogeneous scoping review teams, whose implicit assumptions and epistemological positions or interpretations may also shape the review. Thus, with only 23 (22.7%) author teams describing and reflecting on their team’s characteristics, readers have limited opportunity to consider why and how certain decisions made in the review process may have influenced the review’s conduct, findings, and reporting. Taking a page from qualitative research, we encourage authors to include a brief reflexivity section in which they report and reflect on the characteristics of their team in relation to the study’s design, data collection and analysis, and reporting.^36^ This information increases transparency and allows readers to make informed judgments about the conduct of the review, as well as its findings and interpretations.

The inclusion of external stakeholders in research, including in knowledge syntheses, has been identified as a beneficial component of high-quality, high-impact research.^38,39^ However, in papers reviewed here, only 19.8% of reviews described including external stakeholders. This finding suggests a missed opportunity to improve the execution and usefulness of medical education scoping reviews. To be fair, guidance on which stakeholders to include and how to include them in scoping reviews has been somewhat unclear.^39^ For example, Arksey and O’Malley have only suggested stakeholder inclusion,^7^ whereas Levac has declared it as essential.^10^ On the other hand, stakeholder consultation is absent from the PRISMA-ScR.^13^ Despite this lack of clarity, several scoping review authors appear to be leveraging stakeholders in creative and critical ways. For example, one scoping review, which addressed education to lessen health gaps between Aboriginal and non-Aboriginal peoples, integrated Aboriginal stakeholders throughout the entire conduct of the review.^40^ Undoubtedly, this review would have suffered without stakeholder involvement. As we are unaware of any firm guidance on stakeholder inclusion in scoping reviews within medical education, we propose that an important step forward would be for the field to provide best practice guidelines on the role of stakeholders. Doing so could help to ensure scoping reviews are optimized for medical education.

Nearly half of the included authors chose to conduct a scoping review because of the nature of their topic or the available literature. Specifically, many of these authors commented on the heterogeneity of the literature and its emerging nature such that particular study designs (e.g., randomized controlled trials) were unavailable for review. The ability to include multiple publication types and various materials is often seen as a hallmark of a scoping review. In fact, Arksey and O’Malley declared that “the whole point of scoping the field is to be as comprehensive as possible”.^7^ However, despite their stated rationales, multiple authors limited their inclusion criteria to empirical research and explicitly excluded heterogeneous works such as perspective articles, opinion pieces, and innovations. In so doing, authors may have inadvertently (or advertently) missed work that is important for understanding an emerging research space. Moreover, six authors highlighted the heterogeneity of the included literature as a limitation. We can only guess as to why some authors made the choice to exclude some heterogeneous works (e.g., lack of time, misunderstanding the point of a scoping review, etc.), it does appear there may be some confusion regarding inclusion of various forms of knowledge and/or evidence (a point that has been discussed by Thomas et al., 2019)^4^. To this end, we propose that inclusion and exclusion criteria should be driven by the research question(s). For example, in the current scoping review, we were guided by research questions aimed at understanding the nature of scoping reviews. Thus, we did not include any other publication types because that approach would not allow us to answer our questions of interest.

Twelve reviews described critical appraisal of the articles they included. This contrasts with our finding that in 15 reviews (14.9%), authors cited an inability to conduct an appraisal due to the nature of the scoping review methodology, which they described as a limitation. For example, one author wrote: “The nature of a scoping review eliminates any analysis of the quality of the research conducted, so the information supplied concerning the participants’ comments regarding the usefulness of a peer-coaching approach needs to be interpreted with caution”.^41^ In some cases, authors pointed to the heterogeneity of the literature as a barrier to critical appraisal, but in others there was a sense that in a scoping review critical appraisal is unpermitted. Similar to the inclusion of stakeholders, this appears to be a gray area with limited guidance. To our knowledge there is no specific “rule” that appraisal must or cannot be conducted in a scoping review. That said, the PRISMA-ScR does note that authors should describe critical appraisal of included evidence if critical appraisal is done.^13^ Thus, we encourage researchers to consider their specific review in relation to their research question and the nature of the literature included, and to then make an informed decision about incorporating (or not) clinical appraisal.

Only two scoping reviews were registered and provided links to a submitted protocol.^32,33^ Protocol registration was established to increase transparency in review practices and has been associated with increased review quality.^42^ Additionally, protocol registration can eliminate researchers from embarking on a review already underway. Currently, we are unaware of any journal policies in medical education that require or encourage protocol registration. This begs the question: is it finally time for the medical education community to have a serious conversation about the pros and cons of protocol registration?

### Limitations of this scoping review

We acknowledge a number of limitations in the current study. First, several scoping reviews on other health professions education topics may have been missed because we focused on a core set of medical education journals. Future research should consider expanding the sample of health professions education journals to get a broader view of the field. Second, to guide our data extraction, we used the PRISMA-ScR reporting guidelines, which was published in 2018. It is possible that authors publishing prior to 2018 were unaware of the importance of reporting many of the items in this reporting guideline, and thus did not include them (even if those data had been collected). However, the PRISMA guideline,^31^ which is the basis for the PRISMA-ScR, was published in 2009 and contains the majority of the same items. This suggests that while not specific to scoping reviews, it is possible authors would have had familiarity with the majority of these items.

## Conclusion

In this study, we characterized the scoping reviews published in core medical education journals, including describing the rationales provided for selecting the scoping review methodology. We also examined how authors report the details of their reviews and identified areas for improvement. Taken together, findings from this scoping review of scoping review suggests there is room for improvement in the reporting of scoping reviews, including the alignment of research questions, the creation and publishing of protocols, and the inclusion of external stakeholders in published works.

## Ethical Approval

Reported as not applicable

## Data

The data charting for all studies has been deposited at: https://doi.org/10.6084/m9.figshare.12699698.v1

## Disclaimer

The views expressed in this article are those of the authors and do not necessarily reflect the official policy or position of the Uniformed Services University of the Health Sciences, the Department of Defense, or the U.S. Government.

